# Cost-benefit analysis across smallholder rice farmers reveals that existing fertilization practices severely compromise their income

**DOI:** 10.1101/2024.09.10.612194

**Authors:** Junjiang Chen, Magkdi Mola, Xing Liu, Tien Ming Lee, Nikolaos Monokrousos, Stavros D Veresoglou

## Abstract

Modernizing agricultural practices of smallholder farmers can increase considerably global food produce. Smallholder farmers are, nevertheless, often unwilling to adopt practices unfamiliar to them. Increases in the cost of fertilizers, can nonetheless render existing practices unsustainable, raising concerns of food injustice. We addressed here the potential to which we can increase yield through popularizing existing best practice agricultural approaches. We worked over a network of 40 smallholder farmers in the Danxia Mountain region, Guangdong province, China. We divided them into two crop-rotation treatments and gave them the freedom to implement agricultural management practices the way they usually do. We subsequently clustered them into three groups based on the management history. Over the final growth season, on which the clustering was based, there was an over 30% increase in crop yield between the most and the least efficient clusters. More importantly we show that the use of fertilizers not only did not promote rice production but it also had adverse effects. We finally present a cost-benefit analysis. Through familiarizing of the smallholder farmers’ existing management practices, policy makers could make integrating new farming guidelines easier to adopt than purely dishing out modern farming practice recommendations. We thus propose that any attempts to transition agricultural practices should involve the extensive monitoring of existing practices at the very beginning. This should effectively close existing yield gaps in relation to industrial farmers.

## 1. Introduction

Feeding the ever-increasing global population over the next decades may only be possible if we sustainably intensify agriculture [1]. Smallholder agriculture holds a lot of promise for improving crop produce while conserving the environment [2], but also brings about numerous additional challenges such as coordinating large numbers of farmers and diagnosing suboptimal management practices. Making a more effective use of fertilizers, as an example, can be pivotal in reducing yield gaps and making food production equitable across many small-holder farmer systems [3]. Farmers in China are increasingly encouraged to adopt modern agricultural practices to improve their livelihoods which nevertheless proves tricky to implement [4]. The criteria based on which smallholder farmers decide on management practices occasionally differ from those in intensive farming and could include other than profit, tradition, prosperity, experience, and intuition [2, 4–6]. Young small-holder farmers, for example, often combine city jobs with agricultural activities and can thereby miss the optimal timing for effective management practices (e.g. in-season fertilizer applications) [2, 7]. High labor costs occasionally prevent smallholder farmers from accessing agricultural services for tillage and sowing [5] and lower crop yield. We also know that smallholder farmers tend to use more nitrogen (N) fertilizer than they actually need [4, 8] which can increase the susceptibility of plants to soilborne pathogens [9]. It is consequently probable that some of those practices are compromising crop yield or inadvertently compromise the health of the soil.

Assessing separately the efficiency of the management practices of each individual combination of managment practices across smallholder farms can potentially be daunting due to the large number of small farms. We think, instead, that the way forward is to group management practices into larger classes, describe the classes and assess their relative performance. We could subsequently encourage farmers to follow best-use practices with which they are familiar already, which could effectively address one of the bottlenecks of modernizing agriculture: the reluctance of farmers to adopt cost effective practices with which they are unfamiliar [10, 11]. We feel that the process can speed up the transition towards modern agricultural practices, sustainable grain production, and improve the life-quality of a large number of people (e.g. [12, 13]).

An often cited figure is that smallholder farmers (defined generally as managing farms smaller than 2 ha) contribute to 70–80% of the world’s food [14, 15], but the actual percentage may be less than half of this estimate [16]. According to the latest assessment data, among the 570 million farms worldwide, 475 million are categorized as smallholder farms [17–19], making them collectively a major source of global produce. Specifically, smallholder farmers globally are projected to contribute 30–35% of the world’s food supply, even though they only manage an estimate between 12%-24% of the world’s agricultural land [18, 19]. In China, there are 207.43 million agricultural operating households nationwide, with the vast majority of them comprising farms of less than 1 hectare [4](National Bureau of Statistics of China [20]. One of the primary rice-producing regions in China is the Guangdong province in the South of China [21]. The grain production in the Guangdong Province in 2022 accounted for 12.9 million tons, comprising 11.7 million tons out of which 11.1 million tons, were rice produce (National Bureau of Statistics of China [22]). Guangdong presents one of the most densely populated provinces in China [23], which implies that there is a high demand for ecosystem services to get delivered from agricultural areas.

In this study, we monitored traditional agricultural practices in parallel to farm performance metrics such as crop yield per area and soil properties over a network of 36 farms. We grouped management information across farmers and carried out a cost-benefit analysis per group. We anticipated that the yield gap of the groups that used fertilization practices sparingly compared to those that fertilized heavily would be relatively small and the former groups will have higher net revenue than the latter ones (*Hypothesis One*). This hypothesis builds on the observations of smallholder farmers fertilizing excessively on their properties [4, 8]. We further expected that the efficiency with which groups of farmers use resources varies considerably, meaning that the net revenue also varies considerably across groups (*Hypothesis Two*). In many cases traditional farming practices maintain an astonishing efficiency (e.g. [2] and our aim was to distill management information to identify sets of existing practices that work well and propagate them across smallholder farmers.

## 2. Material and Methods

### 2.1 Site description

The experimental site is located at the Danxia Mountain region (113°36’25’’ N∼113°47’53’’ N, 24°51’48’’ E∼25°04’12’’ E) in Renhua County, Shaoguan City, Guangdong Province, China (Fig. S1). The study area experiences a subtropical monsoon climate with an average annual temperature of 19.6 °C and the average annual sunshine time is 1721 h. The average annual rainfall is 1551.1 mm, with March to August accounting for 75% of the annual rainfall. According to the FAO soil classification, the landform of the experimental site is composed of red glutenite layers and primarily Alfisols [24, 25]. The basic soil physical and chemical properties at 0∼10 cm depth were as follows (means ± standard deviation): bulk density, 1.43 ± 0.01g cm^-3^; pH, 5.26 ± 0.86 g kg^−1^; soil organic carbon, 3.08 ± 1.08 g kg^−1^; soil Olsen-phosphorus, 28.69 ± 13.47 g kg^−1^; soil total nitrogen, 0.20 ± 0.054 g kg^−1^. The content of soil sand, clay and silt is 29.40% ± 12.49%, 22.35% ± 4.66% and 48.25% ± 9.92%, respectively.

### 2.2 Experiment design and management information collection

We set up a network of 40 smallholder farms in September 2022. The network consisted of farms ranging between 200 and 1300 m^2^. Twenty of the farms were located at Xiafu Village, 10 at Chewan Village, and 10 at Xinlian Village (Fig. S1). In each village half of the farms were asked to grow over three growth seasons rice whereas the other half rice with an intermediate fallow period. The growth seasons lasted approximately from April to July for early rice, and July to November for late rice. We monitored management practices from September 2021 till December 2023 (i.e., two late rice and one early rice seasons). The farmers were left to implement (other than crop type) usual agricultural practices which they had to note down and report on those practices weekly. The key information included: (1) Basic information (location details, farmers information); (2) Crop cultivation details (cultivated area, planting time, seed weight, labor duration, labor costs); (3) Fertilization management (fertilizer type, application quantity, cost, application timing); (4) Agricultural chemicals usage (chemicals type, application quantity, usage date, cost); (5) Farm machinery (diesel consumption, usage duration); (6) Yield information (rice yield, selling price). We combined this scheme with extensive monitoring of soil properties and soil health indicators of the farms. Because we initiated the observations after the first growth season had started and we had a lower confidence about the quality of the data we focused on the second and third growth seasons (i.e. one late rice and one early rice seasons). To better compare across farmers, we clustered the farmer practices on the grounds of the practices on the third harvest but report on both the second and the third growth season. The initial experimental design consisted of forty farmers. Over the duration of monitoring, four farmers dropped out of the scheme and we excluded them from further analyses.

### 2.3 Environmental properties

We collected soil samples from all farms in November 2022 and August 2023 at 10 cm depth. After sampling, all soil samples were promptly transported back to the laboratory in insulated boxes with ice packs. Half of the samples were air-dried, sieved, and prepared for the analysis of soil physicochemical properties. The remaining half of the samples were stored in a freezer at -20°C to facilitate further analysis later.

Soil decomposition rates were assayed on December 13 2022 and on September 12 2023 according to Keuskamp et al [26] and we used the tea bag index to summarize values. Soil ammonium nitrogen (NH_4_^+^-N) and nitrate-nitrogen (NO_3_^-^-N) were assayed from soil collected in August 2023. Inorganic N was extracted with a 2 M KCL solution and were analyzed with a Flow Injection Analyzer (FIA star 5000, Germany). Soil pH was assayed on the soil collected in November 2022 and measured at a soil water suspension (soil: 0.01 M CaCl_2_ solution= 1:4) after shaking 30 min with a pH electrode. Soil available P was assayed in soil collected in November 2022, and determined by the phosphomolybdate method [27, 28]. The soil organic carbon (SOC) and total nitrogen (TN) were measured with the potassium dichromate oxidation method [29] and the Kjeldahl method [30], respectively. Soil aggregate stability was measured using a wet sieving apparatus (Eijkelkamp Agrisearch Equipment, Giesbeek, The Netherlands) [31, 32]. Soil particle size fraction was analyzed by a Modified pipette method [33].

### 2.4 Clustering management practices

To maximize compatibility of the data, we worked exclusively on the management data on the third growth season. We considered the following information over that growth season: Total N fertilization, Total P fertilization, Total K fertilization, Number of Fertilization Practices, Total Pesticides applied, Total Herbicides applied, Total Fungicides applied and whether the farmers had been classified in the fallow group or not. To make data comparable, we z-score transformed the variables, meaning that we modified the distribution of each variable so that it had a zero mean and a unit variance. Pesticides, herbicides, and fungicides applied had a smaller effect on crop growth than fertilizer applications: To reduce the weight of these three variables we set the standard deviation to 0.5 (i.e. this was half the variance of the other variables which lowered their relative importance over the clustering algorithm). We also set the two levels categorical variable “fallow group” to the values 0.5 and -0.5. We used k-means clustering to classify farmers into three groups (which was a more parsimonious solution that the four groups we originally tried – based on a within groups sum of squares parsimony criterion.

### 2.5 Cost-benefit analysis

We discriminated across nine sources of costs: N, P and K fertilizer costs, seed costs, tillage and harvest costs, pesticides, fungicides, and herbicides additions. We used median costs per item across all 36 farmers for most costs and cross-checked fertilizer costs with the local supplier. Exceptionally for tillage and harvest the farmers used different services and we used the costs they reported (i.e., ranging between 50 and 385 RMB per mu (1 mu = 0.07 acre) for tillage and 60 to 362 RMB per unit for harvest). Most farmers were using composite fertilizers. To extrapolate the separate costs for N, P and K, we averaged the costs across the prices of several commonly used formulations in the local store. The final values we used were (in Chinese RMB per kg) 13 for N, 30 for P, 10 for K, 83 for fungicides, 100 for pesticides 125 for herbicides and 80 for seeds. We assumed a revenue of 3 RMB per unit rice yield (the revenue from rice was 2.5 RMB/mu at the second harvest and 3.3 RMB/mu for the third harvest; we wanted nonetheless to build our cost-benefit analysis on a fixed revenue per unit yield). The analysis focused on the second and third growth seasons. We analyzed separately the farmers with and without fallow over the second growth season. We assumed a population of 1000 virtual farmers per cluster per fallow group. We then permuted with replacement the usage data within the cluster and fallow group, meaning that each virtual farmers combined observed in the cluster practices. We used this approach to compute the mean costs/benefits and their 95% CIs.

### 2.6 Statistical analysis

To assess the degree to which the three management clusters reflected soil fertility, we engaged into a redundancy analysis. The response variables were the eight variables we used for K-means clustering. The predictors were N, P and K fertilization, rates pesticides, herbicides and fungicide additions, number of fertilization applications and the fallow treatment. We used as predictors –soil organic C, total N, Olsen P, pH, % sand, % clay, NH_4_^+^, NO_3_^-^, Water Stable Aggregates, and the two tea bag indices, TBI K and TBI S.

To address yield relationships across the clusters as well as the degree to which the clusters manifested differences in management practices, we used one-way ANOVA tests with the three clusters as a categorical predictor. In our models assessing rice yield over the third growth season we fitted first the categorical variable “fallow” to account for the fraction of variance that could be explained for the differences in management the preceding growth season. Upon significance (*P*<0.05) of the ANOVA we carried out Tukey post hoc tests to determine for which pairs we had substantial support that they were different. All statistical analyses were carried out on R v 4.2.2 [34] with the help of the library vegan [35] for the redundancy analysis.

## 3. Results

### 3.1 Descriptive statistics

Cluster 3 achieved the highest yield at 448 kg/mu, followed by Cluster 2 with 432.5 kg/mu, and Cluster 1 had the lowest yield at 400 kg/mu (Fig. 1). Cluster 3 had the lowest fertilizer inputs, with respective applications of 8.17 kg/mu, 1.18 kg/mu, and 2.77 kg/mu kg of N, P, and K fertilizer. Cluster 2 applied the highest amounts of P fertilizer (3.04 kg/mu) and K fertilizer (9.13 kg/mu) along with 12.11 kg/mu of N fertilizer. Cluster 1 applied the highest amount of N fertilizer (12.38kg/mu), and P fertilizers and K fertilizers were applied at rates of 2.03 kg/mu and 5.46 kg/mu, respectively. Pesticides were administered at rates (median and in parentheses first and third quartiles) of 574.50 (Q1=361.80, Q3=701.50), 834.50 (Q1=540.00, Q3=1175.80), and 452.22 (Q1=368.20, Q3=799.30) g/mu for

**Fig. 1.**
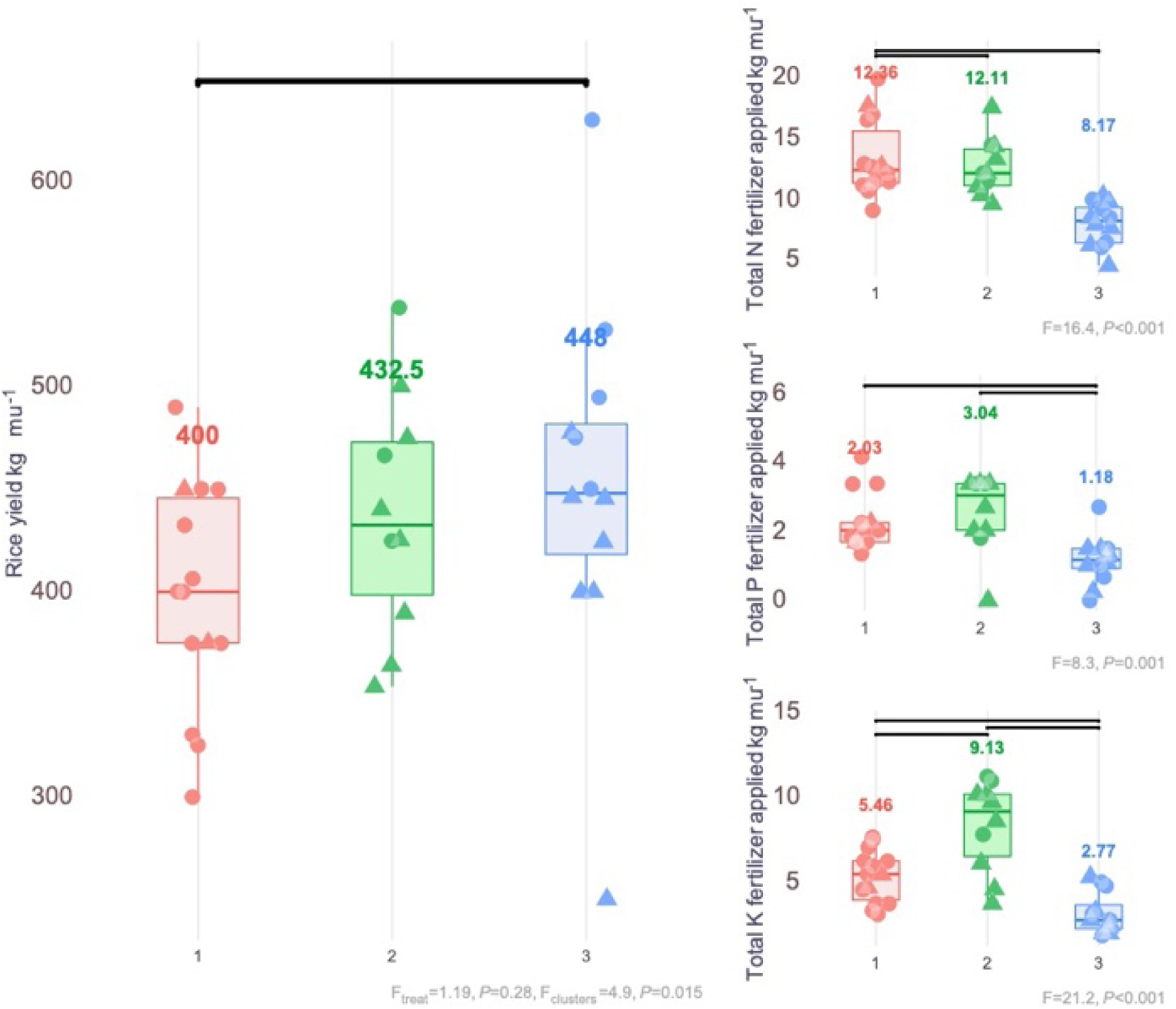
Rice yield (left) and fertilizer application rates of N, P and K (right) across the three management Clusters over the third rice season (which is the season on which the clustering results are based). We used triangles for the farmers that did not undergo a fallow season and circles for those that did. Pairwise significant differences (Tukey Honest significance test) are captured through black horizontal lines. The take home message of the figure is that farmers in Cluster three had a higher yield than those in Cluster one even though they apüplied lower rates of fertilizers.

Cluster 1, Cluster 2, and Cluster 3, respectively. Fungicides were applied to Cluster 1, Cluster 2, and Cluster 3 at rates of 95.50 (Q1=63.49, Q3=139.00), 299.50 (Q1=198.80, Q3=625.80), and 261.94 (Q1=137.30, Q3=357.90) g/mu, respectively. As for herbicides, Cluster 1, Cluster 2, and Cluster 3 received applications at rates of 120.00 (Q1=70.00, Q3=270.00), 70.00 (Q1=60.00, Q3=115.00), and 65.15 (Q1=22.50, Q3=150.00) g/mu, respectively. The median number of fertilization practices across the three clusters were 2.6 (Q1=2, Q3=3), 3.5 (Q1=3, Q3=4) and 2 (Q1=2, Q3=3) respectively for clusters 1, 2, and 3.

### 3.2 Poor relationships between management practices and soil properties

The two first RDA axes collectively explained a total of 28% of variance (Fig. 2). Consideration of additional RDA axes increased this figure but it remained below 40%. The only parameters for which we found support that were related to management practices (ANOVA by terms) were soil texture and soil C. Fertilization practices had positive loading on the main RDA axis, explaining 21.1% of variation, whereas the nutrient availability of N, P and K scored negatively. The three clusters differed (F=9.45, *P*<0.001) on their loadings on this axis with the differences being observed between clusters 1 and 3 (-0.57, *P*=0.018) and clusters 2 and 3 (-0.92, *P*<0.001) (Fig. 2).

**Fig. 2.**
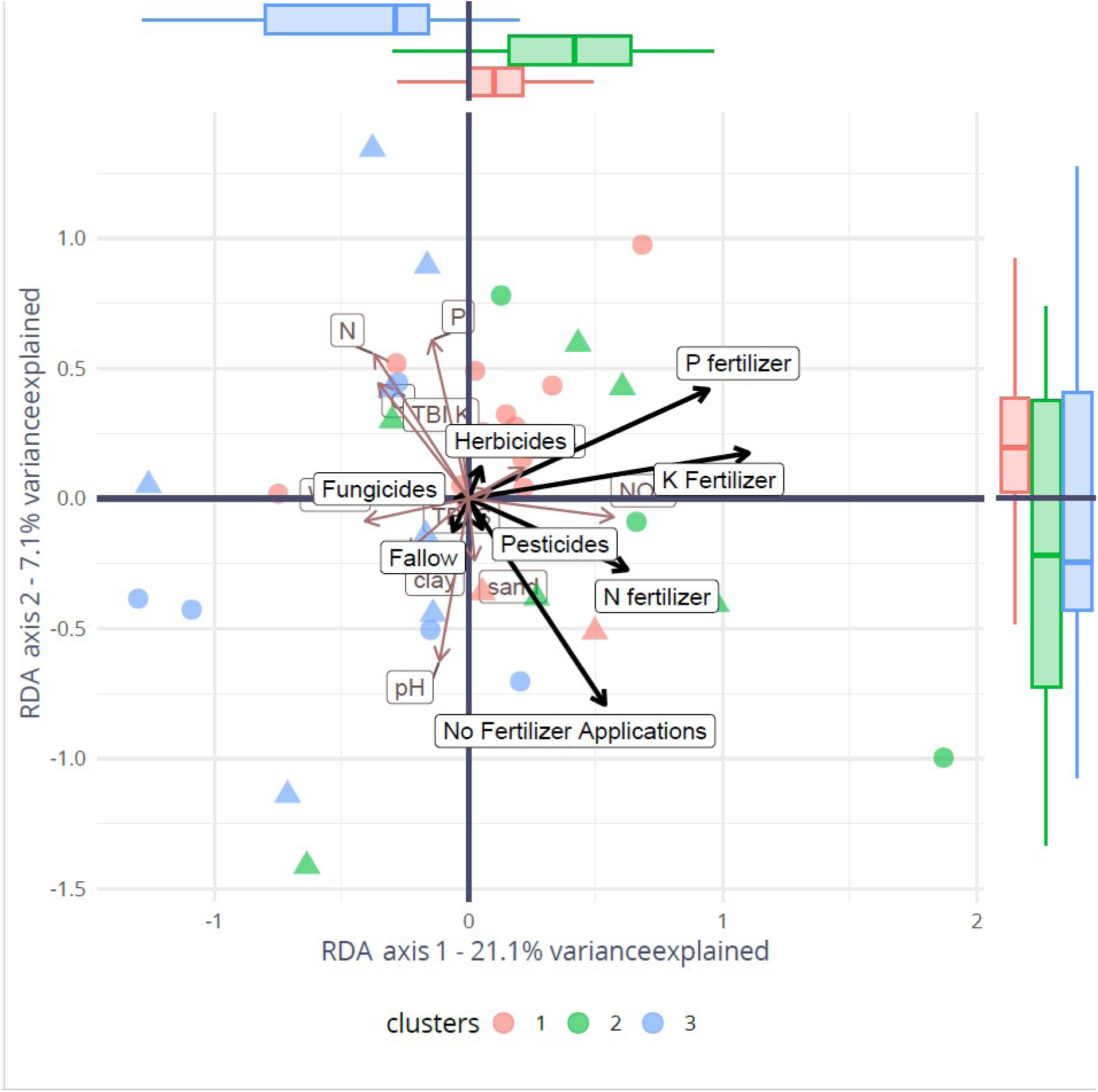
Ordination plot (RDA axes one and two), originating from a redundancy analysis using management practices (black arrows) as response variables and soil properties (red arrows) as predictors. We used circles to highlight farms that were assigned to the fallow treatment and triangles for those in the rice treatment. The take home message of the figure is that there was a poor relationship between management practices and the soil properties in the farm and most soil properties could be collectively summarized over a single RDA axis, RDA axis 1 explaining only a 21.1% of the variance. Note that N, P, K availability in the soil had loadings on the opposite direction over this axis that N, P and K fertilization had.

### 3.3 Rice yield differences across clusters

We observed differences in crop yield (F_clusters_ = 4.9, P=0.015; Fig. 1). We observed a higher mean yield in Cluster 3 than in Cluster 1. Interestingly these two specific clusters differed in the application rates of N, P and K with Cluster 1 having received more of each of these three types of fertilizer. Aside N, P and K fertilization rates (Fig. 1), the three clusters differed in the number of fertilization practices they received (F=13.4, *P*<0.001) with Cluster 2 (median 3.5) differing from Cluster 1 (median 3) and Cluster 3 (median 2).

We observed no differences in yield across the three clusters over the second growth season (Fig. S2). We also observed however that the rates of fertilizer application were inconsistent with those we observed in the third growth season and which had been used to classify the management practices (Fig. S2). Farmers in Cluster 1 applied more N fertilizer than Clusters 2 and 3 (F = 5.8, P<0.016) but there were no differences in P and K fertilizer application rates.

### 3.4 Cost benefit analysis

Net revenue for the farmers in the fallow treatment ranged between 422 (Cluster 1) and 904 (Cluster 3) RMB per mu (Fig. 3a). Differences were due to lower costs but also a higher income across the farmers in the third cluster. The differences were more subtle across the farmers that grew rice over the second season (Fig. 3a) with net revenue ranging between 1092 and 1267 RMB per mu (Fig. 3a). Farmers without a fallow season belonging to the second cluster had the highest expenses but also the highest income from rice. Harvest costs accounted for a considerable fraction of total costs across farmers in the fallow treatment (Fig. 3b). Across the farmers that grew rice over the second season the main cost was the purchase of N fertilizer with considerable resources being spent for seeds and tillage (Fig. 3b).

**Fig. 3.**
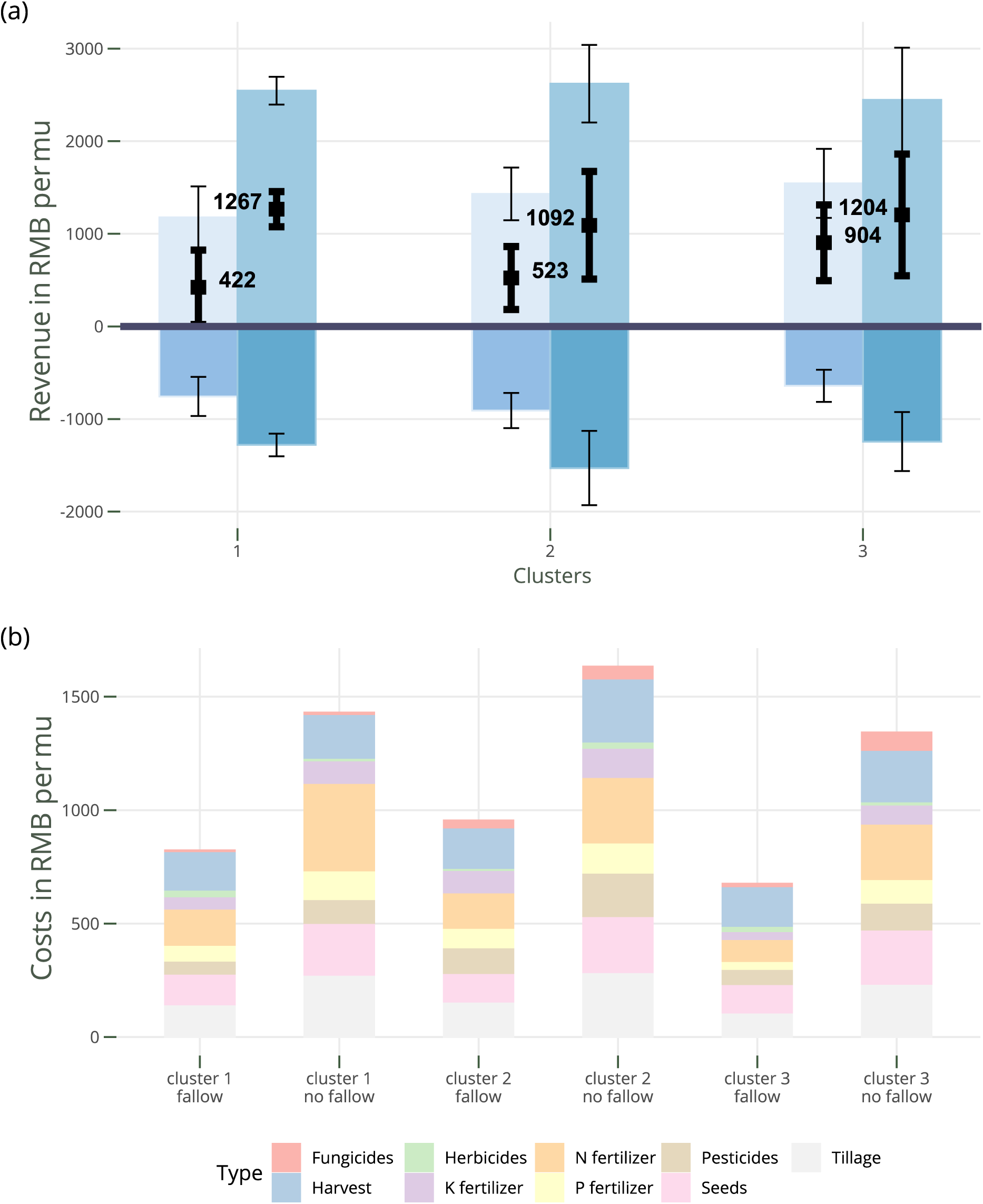
Cost -benefit analysis (a) and cost profiles (b) of the farms across the three clusters of smallholder farmers. Statistics were generated based on a permutation approach. In (a) bars above the zero lines are earnings from crop produce and below the zero line costs of growing rice. Bars on the left in each cluster are for the farmers that were allocated in the fallow treatment whereas those on the right for the farmers that were in the rice season.

## 4. Discussion

We present here a conclusive cost-benefit analysis on how smallholder farmers in Danxia Mountain manage their agricultural properties. We monitored their practices over three growth seasons and we found evidence that fertilizer application rates, not only did they not increase yield, but in many cases, they compromised yield produce (in support with *Hypothesis One*). This compromised considerably the revenue of the farmers that invested in purchasing fertilizers. We also monitored an inconsistency in the way farmers invested in fertilizer applications across growth seasons, which became apparent when we compared the second with the third growth seasons.

Smallholder farmers tend to use N fertilizer ineffectively (e.g. [2, 36], which is exactly what we observed in our study. In the particular case of rice paddies this is undesirable as it cascades to high N_2_O emissions [37] but also aggravates the leaching of N from the fields [38]. Fertilizer application has become an even bigger concern lately because the prices of mineral fertilizers have soared [39, 40]. We informally interviewed farmers about the criteria that determine how much fertilizer they use and they confined to us that their main criterion has been financial availability [41, 42]. They maintain the perspective that they can infinitely improve crop yield through adding fertilizer, which can in many cases lead them to follow the yellow brick road. A staggering 27 of our 36 farms had a pH below 5.5 meaning that the efficiency of mineral fertilizers was low, and even worse mineral N fertilizers could further acidify their fields. It was apparent that the farmers in our study lacked guidance and they had to pay the toll for it. We originally hypothesized (*Hypothesis One*) that the revenue would be higher in the Clusters that used less fertiliyer. We found support for this hypothesis.

There were weak relationships between soil properties, management practices and yield (e.g. Fig. 2). The Technical Guidance for Southern Double-Cropped Late Rice Production in 2019 from the Ministry of Agriculture and Rural Affairs of China [43] recommends applying 5-7 kg of urea per mu 3-5 days after transplanting to promote early growth and an additional 2-3 kg of urea per mu when plants show two leaves inverted and one leaf exposed and 2-3 kg of urea per mu at heading stage. Based on these recommendations, N fertilizer should not exceed 13 kg of urea, whereas the respective recommendations for K and P are 3 kg of KCL, and 200 g of KH_2_PO_4_. These figures correspond to a maximum input of approximately 5.98 kg/mu for N, 0.045 kg/mu for P, and 0.99 kg/mu for K. Farmers in our study (Fig. 1b), greatly exceeded these recommendations, implying that the magnitude to which fertilizers are overused can in some cases be massive (e.g., P in Cluster 2).

The operating profit margin ratios for smallholder farmers is low and it often falls short of meeting their daily household needs. In our analysis we did not explicitly assessed the operating profit margin because of difficulties in assessing interests and worker labor. Cost benefit ratios, however, for the parameters we considered, varied considerably across clusters (Fig. 3a) in agreement with Hypothesis Two. Ren et al. [36], for example, profiled 15000 rural households in China for their income to show that the income they received from agriculture fell severely short of what a young individual would earn as an unskilled worker. An apparent way to slow down the loss (i.e. out-migration) of young Chinese smallholder farmers is through knowledge transfer [44, 45]. Based on our analysis here, simply through transitioning from agricultural practices that we observed in Cluster 1 and Cluster 3 to those of Cluster 2 would secure an improvement in the income by 24.81% and 30.53%, respectively. These figures already exceeded the figure of 22.1% that was reported in the form of a target net gain increase for rice farmers if they implemented modernized farming practices [2]. We thus feel that there is considerable promise in improving smallholder rice growing practices simply through taking advantage of existing knowledge.

We monitored agricultural management practices over a rural area in China and used the data to carry out a cost-benefit analysis. We witnessed some of the issues that smallholder farmers currently face, such as a low operating profit margin and a limited access to farming guidelines, raising concerns of inequalities in food produce compared to industrial farmers. We propose that, at minimum for the early stages of the agricultural modernization transition, proposing alternative management practices could be delivered in the form of assessing the efficiency of existing management practices.

## Supporting information

Dupplementary Display Items

## 5. Declarations

### Funding

The article was funded through the NSFC Research Fund for Outstanding Foreign Young Scholars (grand agreement number 32250610), “Arbuscular mycorrhizae: a land of promise for mitigating terrestrial N_2_O emissions”, awarded to SDV.

### Conflicts of Interest

The authors report there are no competing interests to declare.

### Data and code availability statement

We make the data available in the form of a supplement

### Ethics approval and consent to participate

This study was performed in accordance with the ethical standards as laid down in the 1964 Declaration of Helsinki and its later amendments or comparable ethical standards. The researchers have obtained a support letter from the management committee of the Danxia Mountain Protected Area. Informed consent was obtained from participants involved in this study who were briefed in the presence of a witness (local experts) that their participation in this study was voluntary and that confidentiality would be maintained at all times. All study participants were above 20 years of age and provided informed verbal consent prior to study enrolment. The collected data were kept anonymous, confidential, and in accordance with international and national ethical guidelines. Consent for publication was equally obtained at this point.

### Author Contributions

SDV: Conceived the project; JC carried out the coordination and laboratory work; JC, MM, XL, TML, NM, SDV together wrote the paper, contributed comments and approved the final version of it.

## Notes

### Competing Interest Statement

The authors have declared no competing interest.

