## Supplementary material for "Cost-benefit analysis across smallholder rice farmers reveals that existing fertilization practices severely compromise their income": Dupplementary Display Items

### Supplementary Materials

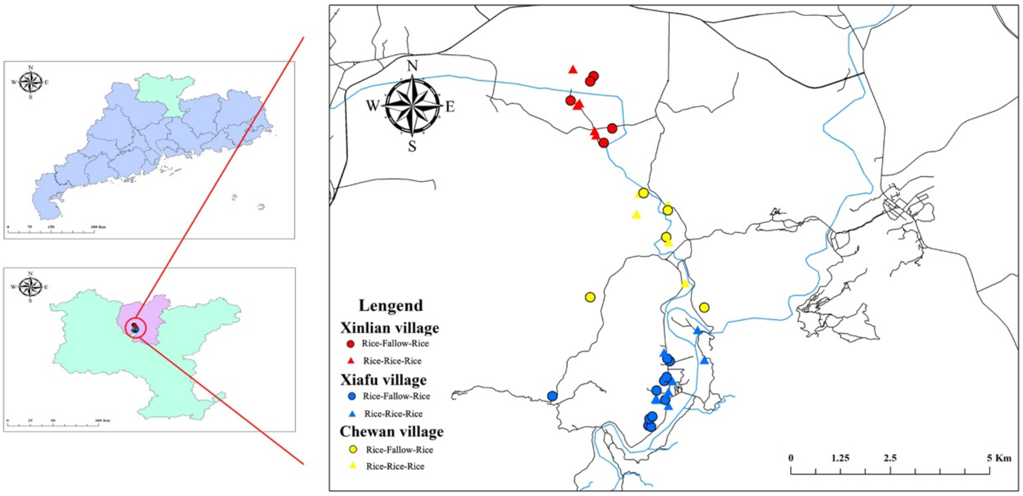


Fig. S1 Location of Danxia Mountain region (left) and distribution of the 40 smallholder farms (close-up map at right). Note: The different colors of the dots represent different villages and different shapes represent different treatments, triangles mean rice-rice-rice treatment, and circles mean rice-fallow-rice treatment.

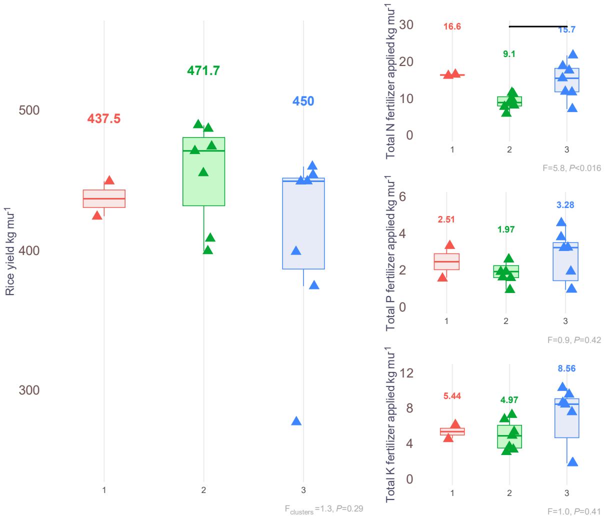


Fig. S2. Rice yields (left) and three different fertilizer application (right) in different groups at second rice season. Note: different colors mean different groups, and different shapes means different treatments. Specially, triangles mean rice-rice-rice treatment and circles mean rice-fallow-rice treatment.
